# intDesc-AbMut: Describing and understanding how antibody mutations impact their environmental interactions^1^

**DOI:** 10.1101/2025.11.18.688974

**Authors:** Shuntaro Chiba, Masateru Ohta, Tsutomu Yamane, Yasushi Okuno, Mitsunori Ikeguchi

## Abstract

Previously, we proposed a double-point mutation (DPM) strategy involving the simultaneous substitution of two amino acids to optimize antibodies. By selecting mutants based on the criterion that favorable interactions between mutated residues and their local environments are preserved or enhanced, improvements in antibody affinity were achieved. Nonetheless, manual extraction of these interactions proved to be time-consuming and labor-intensive. Thus, to streamline this process, we developed intDesc-AbMut, an automated software tool designed to extract interactions between designated mutant residues and their surroundings from three-dimensional antigen–antibody complex structures. intDesc-AbMut identifies and classifies 36 distinct types of interactions, enabling visualization of changes before and after mutation. Additionally, the effects of mutations can be represented by interaction descriptors generated from the extracted interactions. As a practical application, we constructed a machine learning model using these descriptors combined with side-chain rotamer frequencies to determine whether the environment around mutated residues in antigen–antibody complexes is crystal-like, achieving a Matthews correlation coefficient of 0.505 and an accuracy of 0.855. The intDesc-AbMut software is available at: https://github.com/riken-yokohama-AI-drug/intDesc/tree/intDesc-AbMut.

## 1. Introduction

Antibody therapeutics represent a pivotal modality in the pharmaceutical industry, with considerable anticipation surrounding the development of novel antibody drugs [1]. Throughout the antibody development process, structural modifications are essential after a lead antibody is identified to optimize its properties for pharmaceutical use. These modifications focus on improving activity, physicochemical properties, antigenicity-associated toxicity, and pharmacokinetic profiles.

Among the promising approaches to structural antibody modification is single-point mutation (SPM), which involves substituting a single amino acid in the antibody sequence to enhance its properties. SPM strategies utilize the three-dimensional (3D) structures of antigen–antibody complexes [2–4], leveraging both experimentally determined and computationally modeled structures. However, while SPM-based designs have been widely explored, the impact of multiple simultaneous amino acid mutations—particularly those informed by 3D structural data—remains insufficiently investigated. This gap highlights the need for further methodological innovation in antibody engineering.

To broaden the scope of structural modification techniques, we previously introduced a double-point mutation (DPM) strategy that simultaneously mutates two spatially adjacent amino acids within an antibody. Successful application of the DPM strategy has resulted in the generation of multiple antibodies with improved activity [5,6]. Notably, when the two mutations were performed individually as SPMs, the resulting antibodies exhibited markedly reduced activity or were not expressed. This outcome suggests that combining two beneficial SPMs does not necessarily recapitulate the enhanced activity achieved through the DPM strategy.

When developing antibody drugs using the DPM strategy, many antigen/double-point mutant antibody (Ag/DPMAb) complex structures are generated via computational modeling. From this extensive pool of models, those expected to improve key physicochemical properties, such as antibody affinity and stability, must be selected. For example, in a successful DPM application [6], we generated approximately 21,000 Ag/DPMAb complex structural models by introducing 19 types of mutations (excluding cysteine) at each of the two amino acids across 59 residue pairs. MM/GBVI energy [7] values were then used to filter structurally unfavorable models, particularly those exhibiting steric clashes between the two mutated residues. However, MM/GBVI energy was not applied as a criterion for selecting antibodies for further experimental validation, as unfavorable van der Waals (vdW) interactions are readily detectable, while determining whether mutated side chains form favorable interactions is difficult when relying solely on electrostatic and vdW terms, which fail to capture the full spectrum of interactions, such as weak hydrogen bonds (e.g., CH…π and CH…O interactions) [8], orthogonal multipole interactions [9,10], and S…O interactions [11].

Instead, four DPM antibodies were selected by prioritizing mutants with interactions around the mutated residues that were either preserved or enhanced relative to the wild-type (WT) antibody. These interactions were manually identified and analyzed using molecular graphics software; however, this method is time-consuming and introduces uncertainty regarding whether all relevant interactions have been accurately captured.

We aimed to overcome the limitations of manual interaction analysis in this study by developing intDesc-AbMut as a software tool that automatically identifies diverse interactions between mutant residues and their local environments from 3D structural data. It classifies and labels interaction types to aid interpretation, expanding beyond commonly analyzed hydrogen bonds and vdW contacts to include weak hydrogen bonds, S…O, and orthogonal multipole interactions. Visual examination of these interactions is facilitated through molecular graphics. As an application, intDesc-AbMut was used to analyze changes in interactions between WT and mutant antibodies in both SPM and DPM contexts, where previous studies have reported increased binding affinity compared to WT. This tool also generates interaction descriptors that characterize the local interactions surrounding mutant residues. These descriptors detail the types and quantities of interactions present, enabling quantitative tracking of changes in the interaction repertoire upon mutation. This facilitates direct comparison between pre- and post-mutation states. Furthermore, a machine learning model was developed using descriptors of the side-chain structures of specific antibody amino acid residues. The model predicts whether the side-chain structure resembles that observed in the crystal structures, supporting evaluation of whether mutant–antigen complex 3D structures are valid representations. This study examines the performance of this model and the utility of interaction descriptors, with emphasis on identifying which interaction types are most important for classifying crystal-like structures.

## 2. Methods

### 2.1. intDesc-AbMut

intDesc-AbMut is a software tool designed to automatically extract interactions between a designated amino acid residue—intended for mutation—and its local environment from a 3D structure (**Fig. 1**). The environment considered by the tool includes (i) the remainder of the antibody, (ii) the adjacent antigen, and (iii) surrounding water molecules. For each interaction, intDesc-AbMut records the identities of the interacting atom pairs and assigns an interaction-type label (e.g., hydrogen bond, CH…O, etc.). These labels enable the construction of interaction descriptors that enumerate the number of each interaction type present.

**Figure 1.**
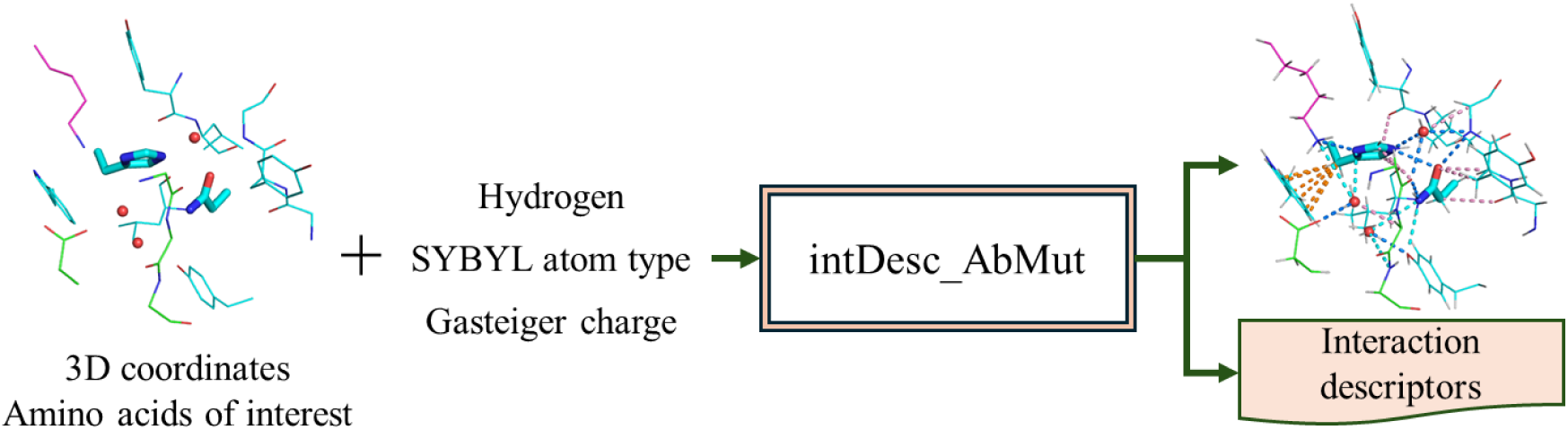
Overview of intDesc_AbMut.

Interactions are primarily defined between heavy atoms, with some exceptions where interactions are defined between bonds. An interaction is assigned only when both the atom types and specific geometric constraints (e.g., distances and angles) are satisfied. Notably, intDesc-AbMut can automatically extract and label 36 different interaction types—including orthogonal multipole, S…O, and weak hydrogen bonds such as CH…O and CH…π interactions. Accordingly, each interaction type has an explicit operational definition. For every heavy atom in the mutant side chain, intDesc-AbMut evaluates the criteria for each interaction and outputs those that meet the requirements.

#### 2.1.1 Definitions of the interactions

Figure 2 illustrates the interaction schemes for CH…O, CH…π, and orthogonal multipolar interactions. The necessary geometric conditions defining each interaction are summarized in **Table 1**. The full set of interaction definitions, including additional interaction types, is provided in **Supplementary Table S1**.

**Figure 2.**
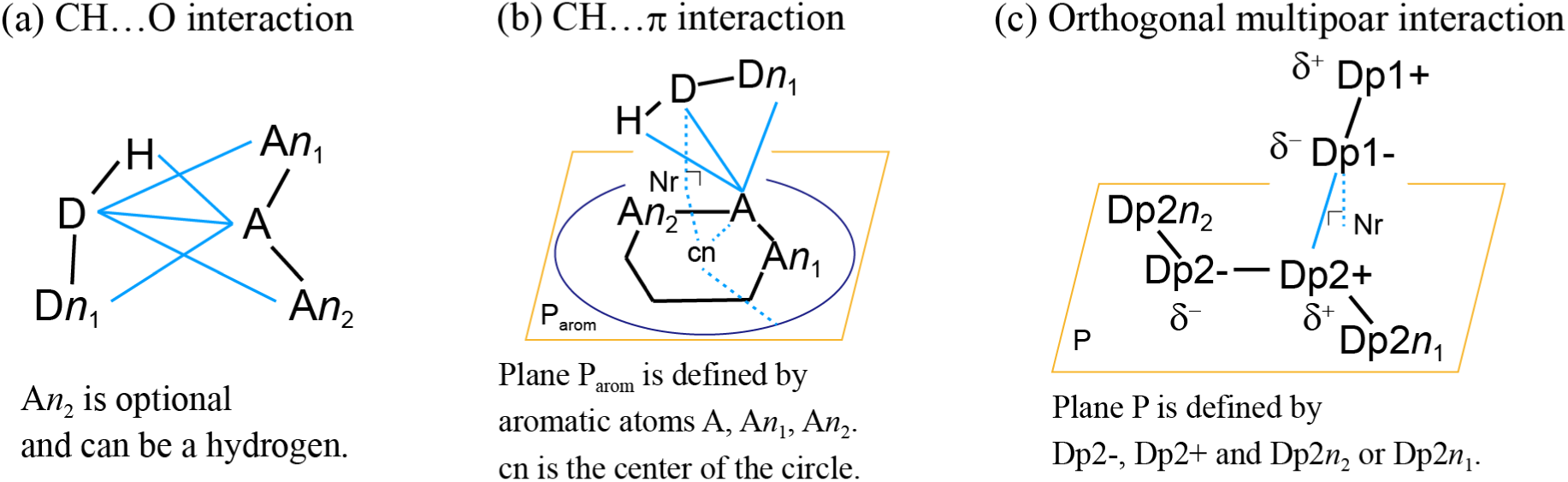
Interaction#schemes of CH…O, CH…π, and orthogonal multipolar interactions. D: Heavy atom attached to donor hydrogen. H: Hydrogen. Dn_x_: x^th^ heavy atom covalently bonded to D. Dpn: Heavy atom of dipole n. Dpn+: Heavy atom of dipole n with a positive partial charge. Dpn-: Heavy atom with a negative partial charge. Black lines: Any type of bond. Cyan lines: Distances between atoms belonging to two different residues. Cyan dashed lines: Dashed cyan line: Auxiliary line for geometric reference. cn: center of the ring composed of aromatic atoms. The circle’s radius is set to X times the cn—A distance (parameter X, default = 1.4) used to evaluate whether a CH…π interaction. P: Plane. P_arom_: plane formed by aromatic atoms. Nr: point where the perpendicular line from the atom to the plane P intersects with P.

**Table 1.**
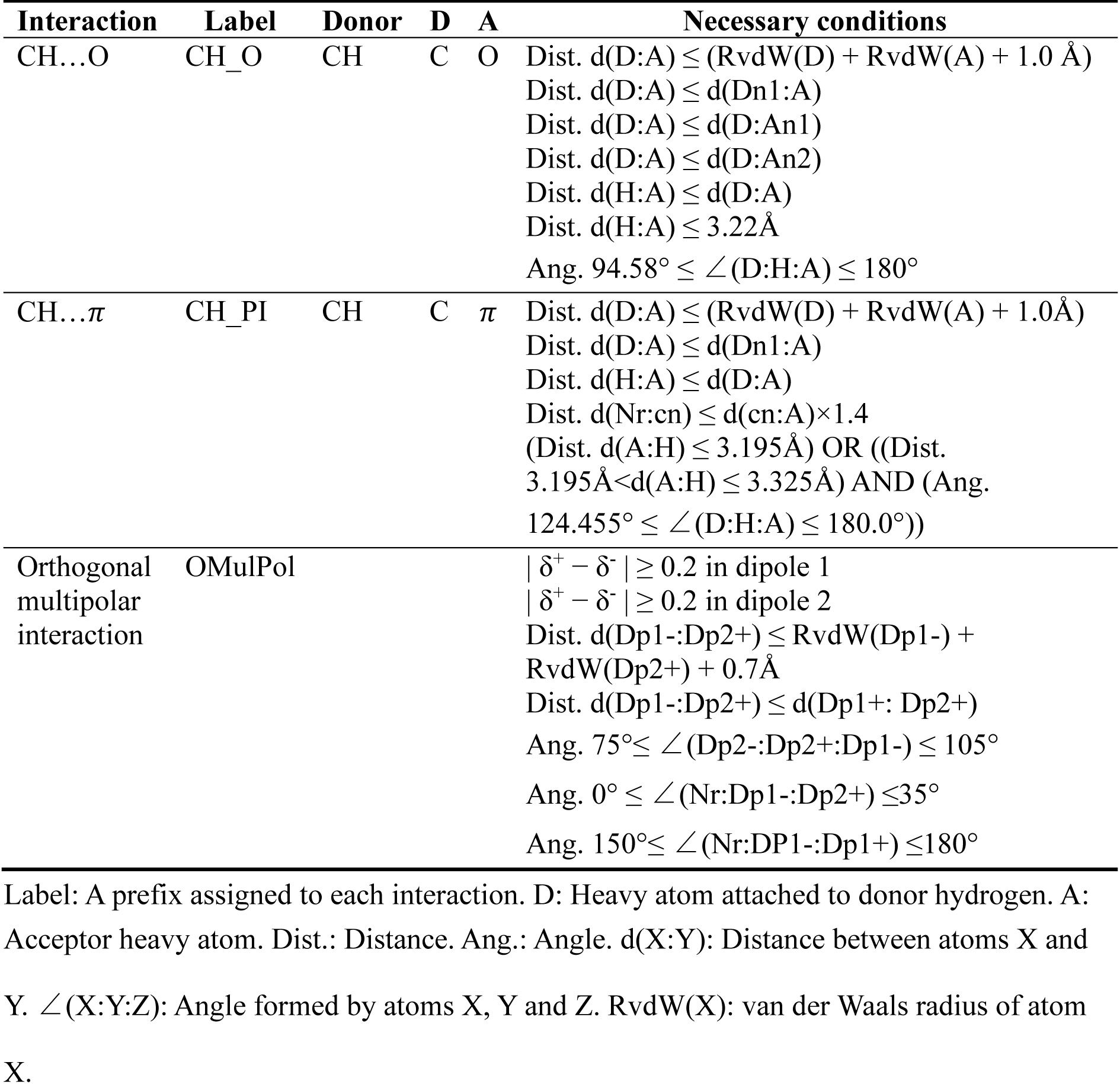
Definitions of CH…O, CH…π, and orthogonal multipolar interactions.

An interaction scheme specifies the motif required for each interaction, encompassing the required atoms, their bonding requirements, atom types (e.g., heavy atoms), definitions of interatomic distances, bond angles, dihedral angles, planes, and, where applicable, constraints on atomic partial charges. In hydrogen-bonding interactions, including weak hydrogen bonds, the acceptor heavy atom is denoted as A, the donor heavy atom bearing the donor hydrogen as D, and the donor hydrogen as H (Figs. 2a and 2b). Atoms covalently bound to D are labeled Dn_1_, Dn_2_, etc., while those adjacent to A are labeled An_1_, An_2_, etc.

For interactions defined between bond dipoles, as in Fig. 2c, the interaction is specified by the atoms constituting each dipole, denoted Dpn+ and Dpn−. Here, the numeral following Dp indexes the dipole, and the subsequent “+” or “−” indicates the atom’s partial charge sign. Likewise, δ^+^ and δ^-^ denote positive and negative partial charges, respectively. The antigen–antibody complex is provided as a MOL2 file that includes per-atom partial charges, enabling atom labeling based on Gasteiger-computed partial charges [12] recorded in the MOL2 file.

**Table 1** details the necessary conditions for classifying each interaction type, where all listed conditions must be satisfied for assignment. Here, “necessary conditions” denote the geometric criteria (e.g., distance and angle thresholds) that must be satisfied for an interaction to be assigned. Constructed points like cn and Nr may replace atom labels where appropriate. All distances and angles are computed from the atomic coordinates provided in the MOL2 file. For dipole-involving cases—such as orthogonal multipolar interactions—the necessary conditions may also include atomic partial-charge specifications (e.g., δ^+^, δ^-^).

Interaction labels comprise the interaction name or abbreviation and may include participating atom names. When visualized in PyMOL, these labels appear as objects and are also written to a text file for programmatic aggregation. By tallying label entries, interaction descriptors can be constructed, such as counting the number of CH…O interactions.

Atom-type requirements for specific interactions (e.g., CH…O) are determined by the SYBL atom types listed in the MOL2 file [13], which encodes element and hybridization (e.g., C.3 for sp³ carbon). intDesc-AbMut determines whether an atom satisfies an interaction’s element criteria by comparing the atom’s SYBYL type with the element specifications in the interaction definition.

The automated extraction of these interactions is applied to the mutated residue and its surrounding residues and water molecules. Surrounding residues may belong to the antibody or antigen. Interactions within the mutated residue are not extracted. Interactions mediated by a single water molecule between the mutated residue and neighboring residues are extracted. **Table 2** presents examples of interactions between a mutated residue (M), antibody (Ab), antigen (Ag), and water molecule (S) extracted by intDesc-AbMut.

**Table 2.** Interaction partners and their descriptor representations.

| Interaction partners | Descriptor representation |
| --- | --- |
| M–Ab | M#Label#Ab |
| M–Ag | M#Label#Ag |
| M–S | M#Label#S |
| M–M* | M#Label#M |
| M–S–Ab | M#Label#S#Label#Ab |
| M–S–Ag | M#Label#S#Label#Ag |
| M–S–M* | M#Label#S#Label#M |

\*Interaction between mutated residues 1 and 2 in case of DPM.
“Label” is a type of interaction label given in **Tables 1 and S1**.

### 2.2. Preparing input structure for intDesc-AbMut

#### 2.2.1 Addition of hydrogen atoms

PDB-format antigen–antibody complex structures typically lack hydrogen atoms required for interaction determination. Hydrogen addition is a required preprocessing step and was performed using the pdb2pqr 2.1.1 program [14,15].

#### 2.2.2 Addition of atom type and bond type information

intDesc-AbMut relies on SYBYL atom and bond types [13], as well as partial charges to determine the types of interacting atoms. Additionally, the input must be a SYBYL MOL2 file with added hydrogens, atomic charges, atom types, and bond types. To facilitate this, the software pdb2mol2.py was developed to convert PDB files (with hydrogens) into MOL2 files that include all necessary information. The SYBYL atom type and SYBYL bond type required for input to intDesc-AbMut can be uniquely determined from the PDB-format amino acid residue name and PDB atom name. This conversion uses a dictionary that maps combinations of amino acid residues and atoms in the PDB file to appropriate SYBYL atom and bond types. The types of amino acids that can be handled by pdb2mol2.py are shown in **Table S2**. PDB files often contain many water molecules, and pdb2mol2.py correctly converts them to MOL2 files.

Since different hydrogen addition programs assign different atom names, distinct dictionary files were created for each program: pdb2pqr 2.1.1 [14,15] and the ff99SB-ildn force field [16,17] assigned by the pdb2gmx module of GROMACS 2021.4 [18]. The desired dictionary file can be specified when running pdb2mol2.py.

Atomic charges of each amino acid are also listed in the dictionary, and partial atomic charge information is added to the MOL2 file accordingly. To calculate atomic partial charges, the N-terminus of each target amino acid is capped with an acetyl group and the C-terminus with NMe, followed by hydrogen addition using AmberTools’ tleap module [19]. Two disulfide-bonded cysteine residues are constructed by capping the N- and C-termini of each residue in a disulfide-bonded state. The amino acid structures are optimized with AmberTools’ Sander module. Gasteiger charges were calculated using the Hgene program (Hydrogen Generation ENginE) [20]. The resulting partial charges are entered into the dictionary for future use.

### 2.3 Files required for execution of intDesc-AbMut

In addition to the MOL2 file, intDesc-AbMut requires several supporting files: an interaction target molecule specification file that identifies the residues of interest, an interaction criteria file with interaction parameter definitions, a van der Waals radius definition file that defines the radius of each element, and an interaction priority file for label assignment when multiple interactions are possible between the same atoms.

### 2.4 Example of application of interaction descriptors generated by intDesc-AbMut

As an example of the application of the interaction descriptors automatically extracted by intDesc-AbMut, we constructed a machine learning model that distinguishes whether each of the multiple side-chain conformations of specific amino acid residues in an antibody is similar to the crystal structure.

#### 2.4.1 Preparing the dataset

A dataset of 520 antibody–protein antigen or antibody–peptide antigen complexes with good resolution (< 2 Å) was obtained from SAbDab (March 8, 2021) [21], a database containing antibody structures from the Protein Data Bank [22]. To exclude crystal lattice interactions, only complexes with a single asymmetric unit were retained. Structures with alternate residue conformations or incomplete side chains were excluded. Interactions cannot be defined when residues have incomplete structures, as their amino acid side chains cannot be determined from electron density. To address this, we selected antigen–antibody complexes with at least one fully determined residue over 4.5 Å from a structurally incomplete residue and a crystallization agent excluded from interaction extraction. This produced 274 PDB entries and complex structures.

#### 2.4.2 Selection of amino acid residues that generate side chain rotamer structures

The machine learning model aimed to predict whether the mutated side chain structure resembles the crystal structure. To generate training data, antibody residues at the antigen–antibody interface with a minimum heavy-atom distance to the antigen of less than 4.5 Å were selected, excluding glycine and alanine, as they lack side-chain rotamers. Additionally, only residues with heavy atoms in the crystal structure that were more than 4.5 Å away from structurally incomplete residues or the crystallization agent were used. These selected amino acid residues, named SPM residues, were used to generate single-residue side-chain structures (Fig. 3a).

**Figure 3.**
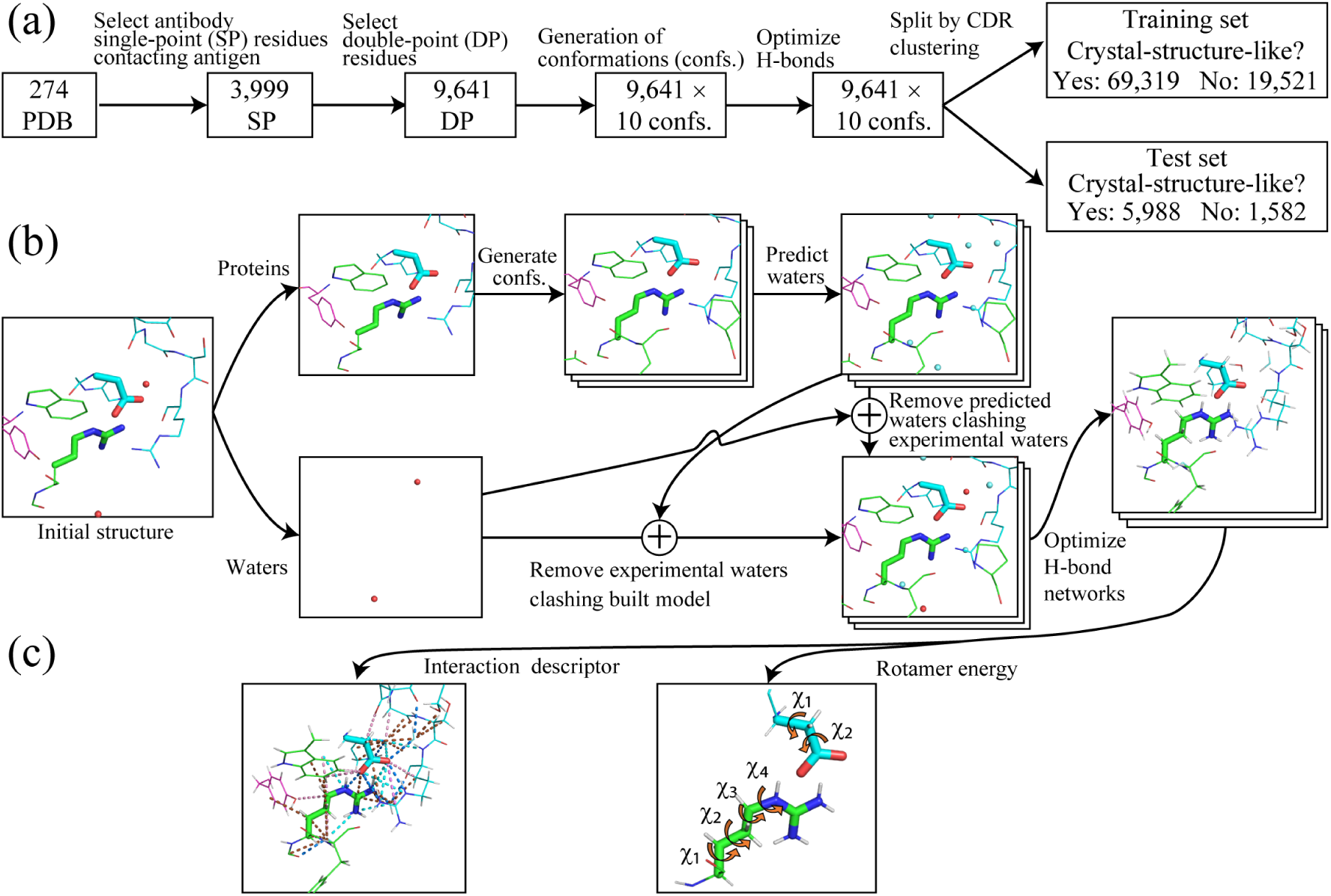
Preparation of training and test datasets. (a) Splitting the training and test sets. (b) Modeling of the side-chain structure and the surrounding water. (c) Descriptor generation.

When the DPM strategy was adopted, two adjacent residues at the antigen–antibody interface were simultaneously mutated, and a 3D model of the two residues was generated. To allow the machine-learning model to assess the likelihood of a DPM structure resembling a crystal structure, a residue pair comprising an SPM residue and another residue within 4.5 Å (minimum heavy-atom distance) was designated as a DPM residue pair. Side chain structures for each DPM residue pair were constructed by combining two residues (Fig. 3a). Glycine and alanine were also excluded from DPM residue pairs.

#### 2.4.3 Generation of side chain structures

For the DPM residue pair, multiple side chain structures were generated. Protein and experimental water molecules were separated, and ten conformations were generated for each mutated residue using FoldX5 [23] (Fig. 3b). Water molecules were docked onto each side chain structure with FoldX4. Experimental water molecules that collided with the docked water molecules (threshold: 1.5 Å inter-heavy atom distance) were deleted; those that did not were returned to their original positions. Hydrogens were then added, and the hydrogen-bond network was optimized using pdb2pqr 2.1.1[14,15].

#### 2.4.4 Determining the crystal structure-likeness of side chain structures

For each generated side chain structure, the root mean square deviation (RMSD) between the model and the crystal structure side chain (excluding *C*_β_) was calculated using the side-chain heavy atom coordinates with the DockRMSD program [24]. Structures with RMSD < 1 Å were labeled “crystal structure-like,” while those with RMSD ≥ 1 Å were labeled “not crystal structure-like.” Symmetry, such as benzene ring rotation, was accounted for in RMSD calculations [24].

#### 2.4.5 Selecting the training and test sets

The training and test sets were selected from the side-chain structure data to maximize antibody diversity, based on complementarity-determining region (CDR) amino acid sequences. First, clustering was performed based on the amino acid sequences of the CDRs for the 274 complex structures (**Figs. S1** and **S2**); the residue numbers of the CDRs were standardized by the IMGT numbering scheme [25] using ANARCI [26] and HMMAR [27] and subsequently clustered into 51 groups using single-linkage clustering implemented in SciPy 1.5.3 [28] (**Figs. S1** and **S2**). The maximum and average CDR identities between cluster members from different groups were 40% and 34 ± 4.9%, respectively. The 51 clusters were randomly split 2:1 into 34 clusters (246 PDB) for the training set and 17 clusters (28 PDB) for the test set. PDB IDs and cluster IDs are provided in **Table S3**. The crystal structure-like and not crystal structure-like data for the training and test sets are shown in Fig. 3a.

#### 2.4.6 Generation and aggregation of interaction descriptors

intDesc-AbMut was applied to all antigen–antibody complex structures in our dataset (Fig. 3a) to extract interactions. Each extracted interaction included the interacting partner and the interaction type, such as M#CH_O#Ab (between mutated residue and antibody residue) or M#CH_O#Ag (between mutated residue and antigen residue). After extraction, interactions were grouped by type and counted, rather than by individual partners, to avoid inflating descriptor counts. For example, in the case of CH…O interactions, M#CH_O#Ab, M#CH_O#Ag, etc., were collectively designated as M#CH_O# (CH…O interaction including mutated residue, regardless of the interacting partner), and the interactions were totaled.

Interactions mediated by water were also grouped and counted as a single descriptor. For example, the number of possible interaction combinations for M#interaction1#S#interaction2#Ab is enormous; thus, rather than counting each interaction combination, they were grouped as an interaction of mutant residues mediated by water, M##S##, and treated as a single descriptor. Similarly, interactions between water molecules and the mutated residue were grouped and counted as a single descriptor. For example, interactions such as M#interaction1#S were grouped as water–mutant interactions (M##S) instead of being counted individually. The five metal interactions and three ion interactions in **Table S1** were excluded from the training data, resulting in 30 interaction descriptors (**Table S4**).

#### 2.4.7 Calculation of side chain rotamer energies

Side-chain structures with high dihedral-angle energies are rare. Additionally, the interaction descriptor is a measure of the goodness of the interaction between the mutated residue and its surroundings, not a direct evaluation of the side chain structure’s poorness. Hence, we developed a rotamer energy calculation program, rotamer_frequency.py, to apply rotamer energy as a descriptor of structural “poorness,” as calculated using the following equation:

*Energy = -RT ln(probability/maximum probability)* where *R* and *T* are the gas constant 0.001987 kcal/(mol K) and the temperature 300 K, respectively.

Rotamer probabilities for residue conformations were obtained from a backbone-dependent rotamer library that compiles the relationships between rotamer conformations and the frequencies of experimental structures [29]. The maximum value of each rotamer probability for a side chain was selected from the probability values defined for the same backbone geometry. The rotamer probabilities of the two amino acid residues constituting DPM were assumed to be independent and calculated by multiplying the existence probabilities of each residue.

#### 2.4.8 Model training

A learning model was constructed using 30 interaction descriptors to classify whether a side-chain structure is crystal structure-like (RMSD < 1 Å). The training set comprised 34 CDR clusters. XGBoost 1.5.1 [30] was used as the machine learning method, with hyperparameters optimized by grid search via Scikit-Learn 1.5.0 [31] (**Table S5a)**. The group shuffle partitioning method was applied, using 64% of clusters for model building, 16% for monitoring model performance (early stopping), and 20% for evaluating built models. The CDR cluster groups defined in section 2.4.5 were used to divide the group units. Boosting rounds were capped at 20,000, with early stopping applied if no improvement in the Matthews correlation coefficient (MCC) [32] was observed after 100 consecutive rounds. The learning rate was set to 0.025. Grid search was repeated 200 times, and the hyperparameter maximizing the average MCC was selected. With selected hyperparameters, 80% of clusters were randomly used for training and 20% for early stopping (**Table S5b).**

A second machine learning model was created using 31 descriptors: 30 interaction descriptors plus the rotamer energy descriptor (**Table S5b)**.

#### 2.4.9 Contribution of each descriptor in the machine learning model

The contribution of each descriptor to improving the predictive accuracy of the machine learning model was evaluated using the permutation importance method [33]. The MCC calculated on the test set using all descriptors was defined as the baseline performance.

Subsequently, the values for each descriptor were randomly shuffled, and the MCC was recalculated 100 times. The mean reduction from baseline MCC was used as the significance criterion.

#### 2.4.10 Calculation descriptor contributions to crystal structure-likeness classification

To assess the ability of a model to predict crystal structure-likeness based on the values of each descriptor given as inputs for a given side chain structure, SHAP values (SHapley Additive exPlanations) [34] and their implementation (https://github.com/shap/) were used. SHAP values, based on game theory, quantify the additive contribution of each descriptor to the prediction.

## 3 Results and Discussion

### 3.1 Application and visualization of intDesc-AbMut

To demonstrate the application of intDesc-AbMut, we examined the SP mutant H-N57Y in the Tissue Factor–anti-Tissue Factor antibody complex, where Asn57 in the heavy chain of crystal structure [35] was mutated to Tyr. This mutation increased the binding affinity of the anti-Tissue Factor antibody by 3.1-fold [6]. The WT structure was generated by adding hydrogens to the crystal structure (PDB ID: 1JPS) using pdb2pqr, followed by conversion to a MOL2 file with pdb2mol2.py. The mutant H57Y model was constructed in our previous study [6], in which hydrogens were added to the crystal structure (1JPS) using Protonate3D [36] in MOE [37], and the structure was optimized using the ff14SB molecular force field. Mutant structures were created using MOE’s residue scan module, output as PDB files, and converted to MOL2 files using pdb2mol2.py.

#### 3.1.1 Interactions of the WT (N57)

Using the MOL2 files for the WT and mutant, intDesc-AbMut was employed to automatically determine interactions between residue 57 and its surroundings in both structures, producing a PyMOL Script file (pml file) for visualization. Loading the corresponding MOL2 file as input for intDesc-AbMut into PyMOL [38] and the pml file enabled clear visualization of these interactions (Fig. 4).

**Figure 4.**
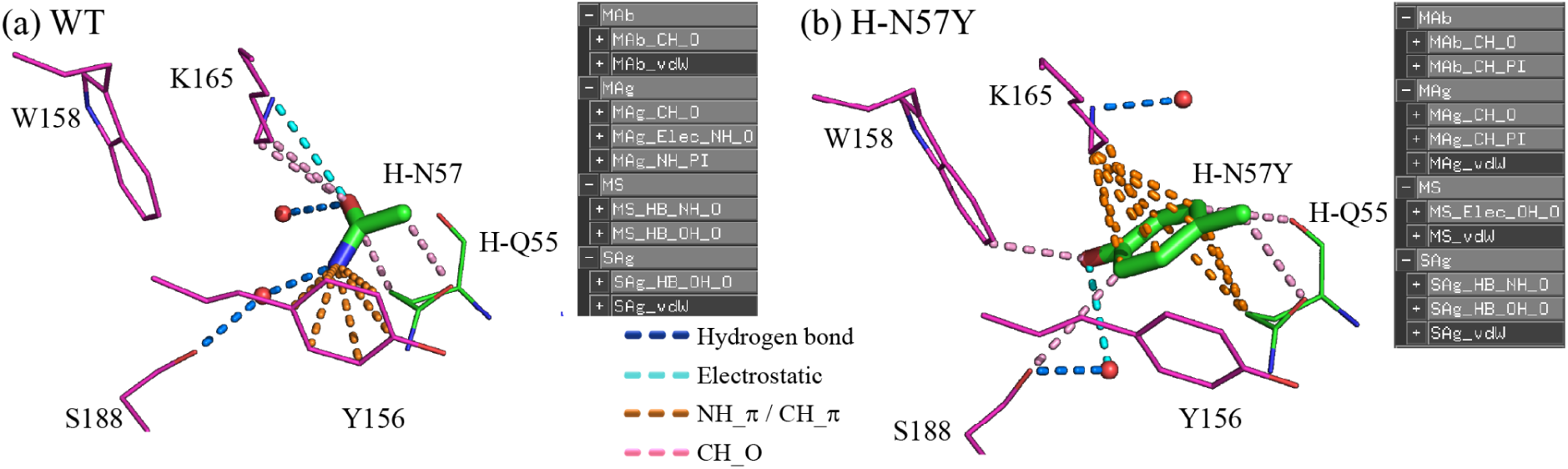
Changes in interactions caused by the H-N57Y SP mutation. (a) The wild type. (b) The SP mutant H-N57Y. Hydrogen bonds are shown in blue, electrostatic interactions in cyan, NH…π and CH…π interactions in orange, and CH…O interactions in pink. van der Waals (vdW) interactions are not shown. Green indicates antibody, and magenta indicates antigen. Hydrogens are not shown. Only the side chain structures of N57 and N57Y are shown in stick form. Only the oxygen atoms of water molecules are shown as red spheres. W158 in (a) does not interact with the side chain, but is shown for comparison with (b).

Analysis of the WT heavy chain Asn57 (H-N57) side chain with intDesc-AbMut revealed numerous NH…π interactions with Y156 (Fig. 4a). In this interaction, the hydrogen atom of NH acts as a donor and the π electrons of the aromatic ring act as an acceptor. The interaction between the NH_2_ group and S188 was mediated by water through hydrogen bonds. The carbonyl oxygen of the H-N57 side chain, acting as an acceptor, interacted with the *N*_ζ_H_3_ of the K165 side chain, acting as a donor. Electrostatic interaction, MAg_Elec_NH_O, was assigned instead of a hydrogen bond because the distance between N and O was greater than 3.2 Å. intDesc-AbMut distinguishes hydrogen bonds (≤ 3.2 Å) and electrostatic interactions (> 3.2 Å) based on interatomic distances, and visualizes hydrogen bond distances with color changes. The carbonyl oxygen of the H-N57 side chain also formed CH…O interactions with the *C*_δ_H_2_ and *C*_ε_H_2_ of the K165 side chain and the *C*_γ_H_2_ of the antibody H-Q55 side chain, indicating weak hydrogen bonds, in which the hydrogen of CH acted as a donor and the O as an acceptor. Thus, carbonyl oxygen recognition of the H-N57 side chain was achieved through electrostatic interactions with the *N*_ζ_H_3_ of K165 and through weak hydrogen bonding with the CH_2_ side chain. Additionally, *C*_β_H_2_ of the H-N57 side chain interacted with the carbonyl oxygen of the antibody H-Q55 side chain via CH…O.

#### 3.1.2 Interactions of the N57Y mutant

In the H-N57Y SP mutant model, intDesc-AbMut revealed loss of the NH…π interaction with Y156 and the CH…O interaction between the *C*_δ_H_2_ and *C*_ε_H_2_ of the K165 side chain and the carbonyl oxygen of the N57 side chain that were observed in H-N57 (Fig. 4b). However, the phenol ring of the H-N57Y side chain, introduced by mutation, formed new CH…π interactions with the *C*_δ_H_2_ and *C*_ε_H_2_ of the K165 side chain and the *C*_γ_H_2_ of the antibody H-Q55 side chain, surpassing the lost interactions. The water-mediated interaction with S188 shifted from NH2 in N57 to phenol OH in N57Y, and a CH…O interaction between the N57Y side chain *C*_ε_H and the S188 side chain was added. W158 of the antigen, previously non-interacting, formed a CH…O interaction with the side chain indole ring *C*_η_H as the donor and the phenolic oxygen atom of N57Y as the acceptor. These interaction changes accounted for the increased activity of H-N57Y, consistent with previous studies. [6]

#### 3.1.3 Interactions of the WT (N34/H91)

A second example involved intDesc-AbMut analysis of the L-N34D/H91S mutant, which exhibited increased binding affinity from 45 pM to 25 pM. The WT structure was generated using methods similar to those for the H-N57Y mutant. The L-N34D/H91S mutant structure was similarly created based on the 1JPS crystal structure using MOE’s residue scan.

In the WT, a network of hydrogen bonds formed between the antigen K169 and the antibody H91, between H91 and N34, and between N34 and Y50 (Fig. 5a). This structure was reinforced by CH…π interactions between the C_β_ of H91 and W96, CH…O interactions between the C_δ_H of H91 and both L89 and Y32, and CH…O interactions between the O_δ_ of N34 and Y49. The N_ε_H of H91 formed a hydrogen bond with water, facilitating additional hydrogen bond and CH…O interactions with Y50.

**Figure 5.**
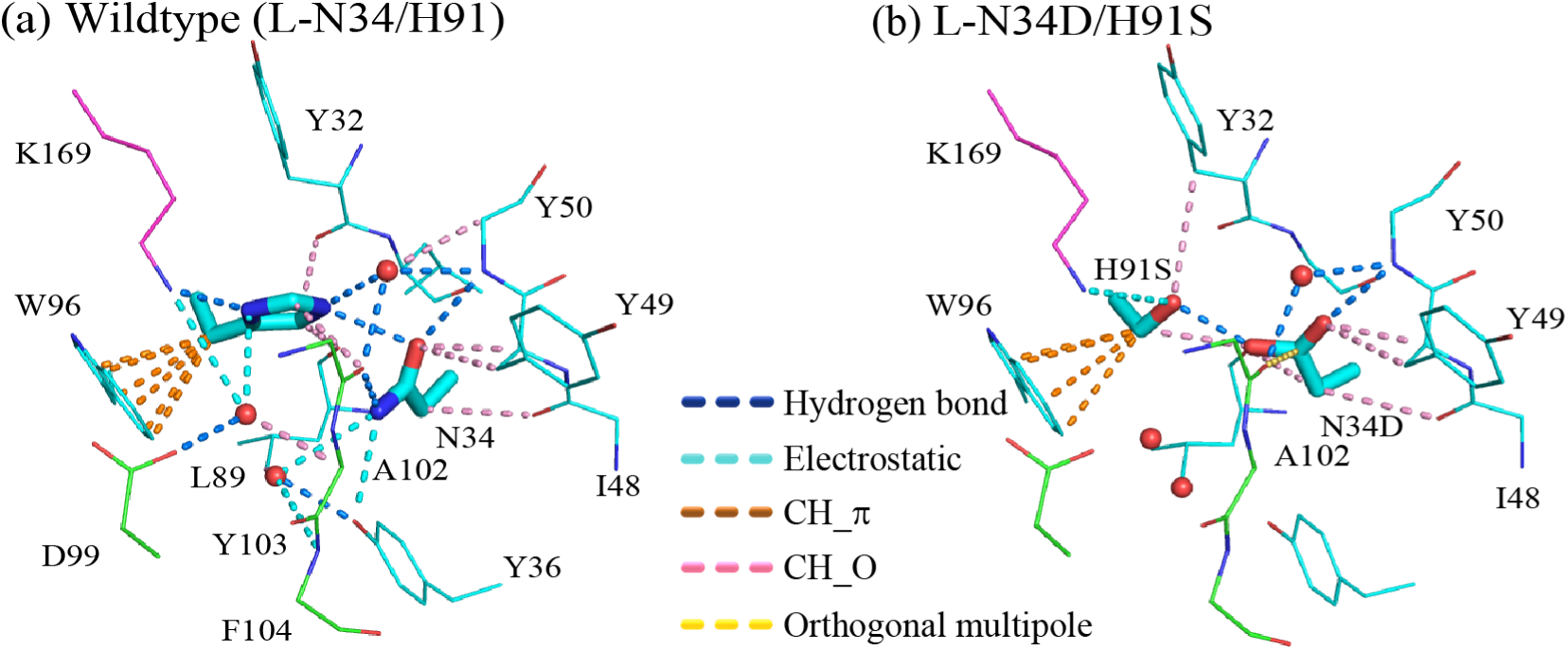
Changes in interactions caused by the L-N34D/H91S DP mutation. (a) The wild type. (b) The DP mutant L-N34D/H91S. Hydrogen bonds are shown in blue, electrostatic interactions in cyan, CH…π interactions in orange, CH…O interactions in pink, and orthogonal multipole interactions in yellow. van der Waals (vdW) interactions are not shown. Green indicates antibody heavy chain, cyan indicates antibody light chain, and magenta indicates antigen. Hydrogens are not shown. Only the side chain structures of N34, N34D, H91, and H91S are shown in stick form. Only the oxygen atoms of water molecules are shown as red spheres. No interactions with Y36, L89, D99, or F104 are shown in (b), but are displayed for comparison with (a).

#### 3.1.4 Interactions of the N34D/H91S mutant

For the L-N34D/H91S mutant, the K169–H91S interaction was classified as electrostatic due to a 3.9 Å distance, but remained present (Fig. 5b). Additionally, hydrogen bonds formed between H91S and N34D, and between N34D and Y50, creating an interaction network similar to that of the WT. This network was further reinforced by CH…π interactions between the C_β_ of H91S and W96, CH…O interactions between the C_γ_ of H91S and Y32, CH…O interactions between the C_β_ of H91S and one of the O_δ_ atoms of N34D, and CH…O interactions between the other O_δ_ atom of N34D and Y49. The backbone carbonyl oxygen of L89 formed a CH…O interaction with the C_δ_H of H91 in the WT, whereas in the L-N34D/H91S mutant, it formed a CH…O interaction with the C_β_H of N34D. Additionally, an orthogonal multipolar interaction between the backbone carbonyl oxygen of the antibody heavy chain A102 and the C_γ_ of N34D was detected in the L-N34D/H91S mutant but was absent in the WT. Orthogonal multipolar interactions occur when the δ⁻ of a dipole, such as C=O, interacts perpendicularly with the δ⁺ of another dipole. Overall, intDesc-AbMut demonstrated that the interactions in the WT were preserved or enhanced in the L-N34D/H91S mutant, supporting its selection for experimental validation.

The ability to visually and easily define the types and numbers of interactions between specific residues and their surroundings in WT and mutant structures is valuable for understanding mutation effects.

### 3.2 Interaction descriptors as machine learning features

From this point forward, we present the use of interaction descriptors in a machine learning model that evaluates whether the side-chain structure of a specific residue or residue pair in an antigen–antibody complex is crystal structure-like. The confusion matrix and statistical metrics using only the 30 interaction descriptors are summarized in **Table S6a**. The test set comprised 5,988 “crystal structure-like” and 1,582 “non-crystal structure-like” data points derived from 28 antibody–antigen complex structures. The model correctly classified 5,730 and 449 of these, respectively. The overall classification accuracy was approximately 0.816, while the MCC was 0.336. Given that the test structures originated from antibodies with different CDR sequences, the results suggest that interaction descriptors can, to some extent, determine whether a side-chain model structure resembles a crystal structure.

While an MCC of 0.336 is not poor, it does not reflect strong model performance. We hypothesized that interaction descriptors primarily capture favorable interactions, but do not directly represent unfavorable side-chain conformations. Accordingly, we incorporated a rotamer energy term into the descriptor, penalizing unfavorable side-chain conformations. Reconstructing the model with the rotamer energy term improved the MCC to 0.505 and the accuracy to 0.855. This suggests that evaluating favorable interactions and penalties for unfavorable rotamers enhances the determination of crystal structure-likeness. As evident from the confusion matrix, adding rotamer energy increased the correct classification of crystal-like and non-crystal-like side-chain structures (**Table S6b**).

### 3.3 Contribution of each descriptor to side chain structure discrimination

The permutation importance method [33] was employed to determine the contribution of each descriptor to the predictive accuracy of the machine-learning model (Fig. 6). The principal descriptors for judging whether a side-chain conformation is crystal-like are the rotamer energy (Rot_energy), CH…π interactions (M#CH_PI#), vdWs interactions (M#vdW#), and CH…O interactions (M#CH_O#). Hydrogen bonds (M#HB_NH_O and M#HB_OH_O) are also important, though to a lesser extent.

**Figure 6.**
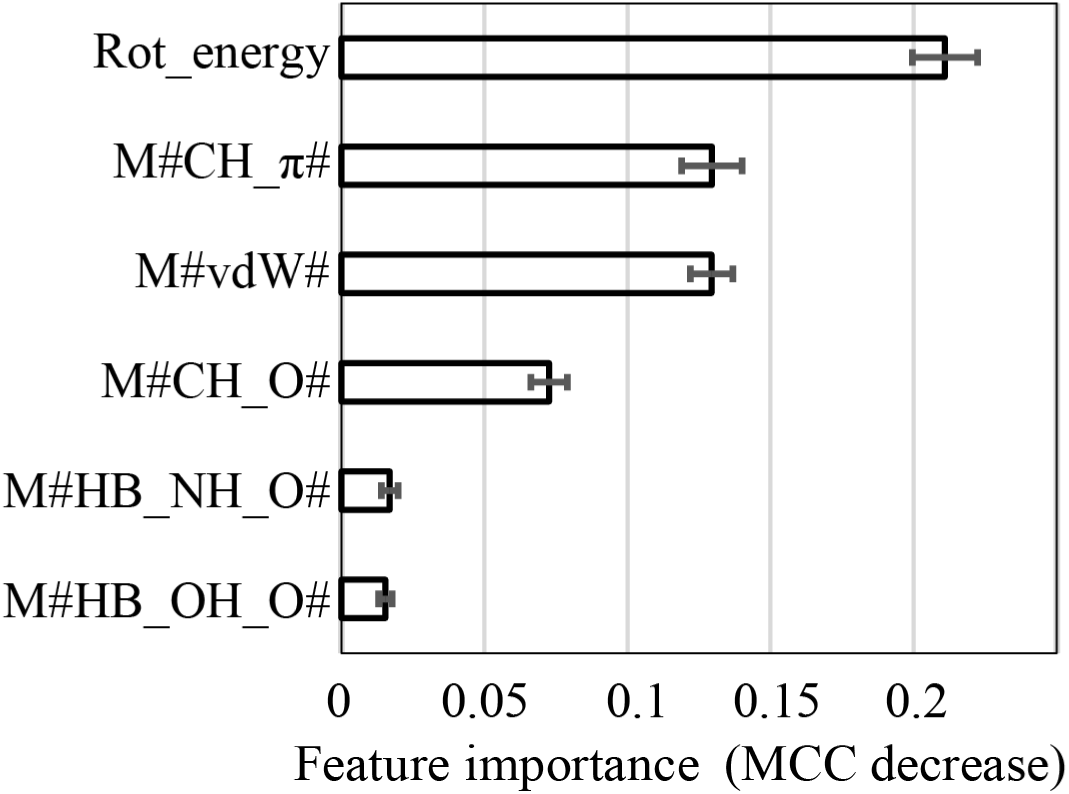
Importance of each descriptor. Error bars represent standard deviations calculated from 100-time repeated evaluations. Only the six top contributing descriptors are listed. The complete list of importance is shown in Table S4.

The rotamer-energy term reflects the occurrence probability of side-chain rotamers; lower-probability conformations are penalized, thereby influencing similarity to crystallographic structures. CH…π interactions represent weak, dispersion-driven hydrogen bonds between the aromatic ring and the neighboring C–H group. Aromatic amino acids such as Phe, Tyr, Trp, and His tend to maximize CH…π interactions in crystal-like structures, whereas in incorrectly modeled conformations, CH…π interactions are not efficiently formed, reflecting this difference.

In our definitions, vdW interactions comprise those between neighboring heavy atoms not assigned to other interaction types. Thus, vdW interactions exclude electrostatically favorable interactions—including weak hydrogen bonds—and favorable dipole–dipole interactions. Hence, they primarily reflect steric shape complementarity and packing between the mutant residue and its environment. It is therefore reasonable that superior geometric complementarity would promote a more crystal-like side-chain conformation.

CH…O interactions, also weak hydrogen bonds, contribute significantly to crystal-like side-chain conformations, as proteins are rich in oxygen atoms—including backbone carbonyl oxygens and those in the side chains of Ser, Thr, Asn, Gln, Asp, and Glu—and donor hydrogens, such as backbone C_α_–H and side-chain C–H, CH₂, and CH₃ groups. The high polarity of oxygen atoms allows them to participate in strong (NH…O and OH…O) and weak (CH…O) hydrogen bonds, thereby stabilizing crystal structures.

Notably, CH…π and CH…O interactions, as weak hydrogen bonds, play significant roles in determining the crystal structure-likeness of side-chains, emphasizing the value of intDesc-AbMut for visual inspection and understanding their formation.

### 3.4 Examples of successful machine learning model predictions and descriptor contributions

Two examples of modeled structures for the residue pair Tyr33 and Gln35 (DPM) in the antibody light chain (PDB ID 6VJN[39]) were evaluated (Fig. 7). The machine learning model, using both interaction descriptors and rotamer energy, successfully distinguished the correct side-chain structure (Fig. 7a) from the incorrect side-chain structure (Fig. 7b). The “correct” modeled structure was predicted to be crystal structure-like with a probability of 0.856 and a side-chain RMSD of 0.2 Å relative to the crystal structure. In contrast, the “incorrect” modeled structure had a predicted probability of 0.186 and a significantly higher RMSD of 6 Å.

**Figure 7.**
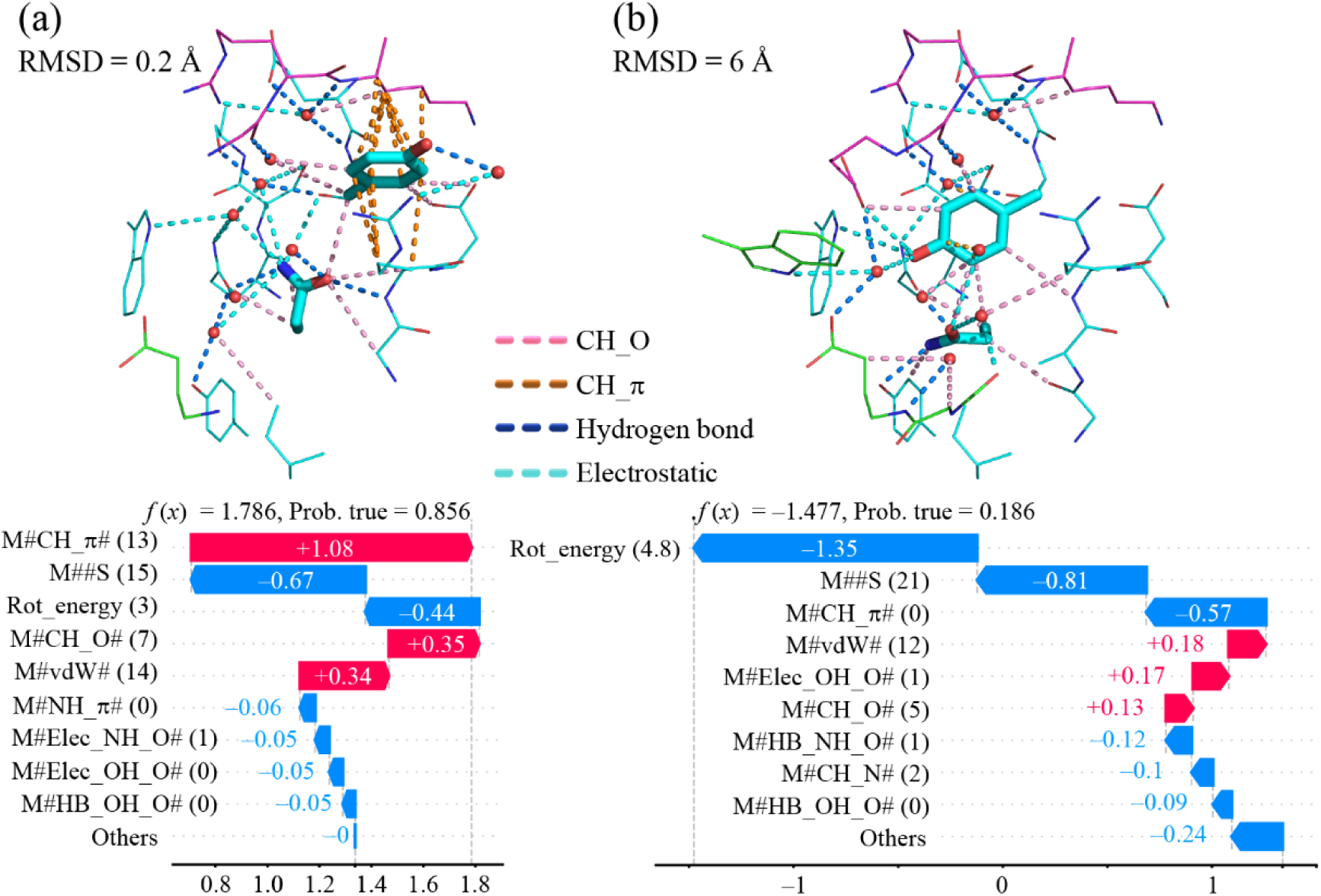
Contribution of interaction descriptors to the prediction value in the side-chain models. (a) Modeled structure correctly predicted as “crystal structure-like.” (b) Modeled structure correctly predicted as “non-crystal structure-like.” Tyr33 and Gln35 from PDB ID = 6VJN [39] were used. Cyan: antibody light chain; green: antibody heavy chain; magenta: antigen. vdW (van der Waals) interactions are not shown. Hydrogen atoms are not shown. Only the side-chain structures of Y33 and Q35 are shown in stick form. In water molecules, only oxygen atoms are shown as red spheres. The contribution of each feature to the probability of crystal structure-likeness (Prob. true) was calculated using SHAP values [34], indicated as *f* (*x*).

Inspection of the SHAP values for the case correctly predicted as crystal-like showed that weak hydrogen bonds, specifically CH…π and CH…O interactions, contributed positively to the classification, followed by vdW interactions, reflecting steric shape complementarity (Fig. 7a). In contrast, for the case predicted as not crystal-like, the positive contributors were vdW, electrostatic OH…O interactions, and CH…O interactions, all of which exerted small effects (Fig. 7b). The strongest negative contributors to the crystal-like classification were the rotamer-energy term, water–side-chain interactions, and CH…π interactions.

## 4 Conclusion

In our previous work, we successfully selected DPMs with improved binding affinity from numerous 3D structural models by evaluating whether interactions between the mutated amino acids and their local environments were maintained or enhanced relative to the WT [6]. Building on these results, we developed intDesc-AbMut to automate interaction extraction, which had previously been performed manually. intDesc-AbMut visualization simplifies the interpretation of mutation effects as changes in the underlying interaction network.

Furthermore, intDesc-AbMut labels diverse interaction types and generates descriptors summarizing the interaction state between mutant residues and their local environments. In a machine-learning classifier tasked with determining whether a modeled side-chain conformation was crystal-structure-like, these interaction descriptors constituted highly informative predictive features.

Despite these advances, neither intDesc-AbMut nor its descriptors alone enabled automatic selection of mutants with improved binding affinity, primarily due to the limited number of practical DPM implementations—only four DPMs were experimentally characterized, with two showing enhanced affinity. For the tens of thousands of modeled DPMs that have not been experimentally tested, improved affinity remains unknown, necessitating additional data before the proposed approach can be rigorously evaluated as a means to automate DPM candidate selection.

The machine-learning classifier was designed and trained specifically on crystal structures or protein or peptide antigen–antibody complexes, thus its applicability is limited to side-chain rotamers within such complexes. Moreover, the model assumes the backbone remains unchanged from its crystallographic coordinates, making it unsuitable for structural alterations such as insertions or deletions. With these constraints in mind, the model may prove useful in combination with structure-quality metrics such as GA341 [40], ProQ [41], or MolProbity [42].

The automatic extraction, visualization, and description functions developed in this study can also be applied to interactions between small-molecule ligands and surrounding proteins. Related software has been developed, with future reports planned. However, while the current analyses were conducted using static structures, applying this approach to molecular dynamics trajectories could enable tracking of dynamic interaction changes over time; the corresponding software has also been developed and will be reported in the near future.

Overall, this approach extends and supports the DPM strategy, aiming to make DPM a practical and accessible option for antibody optimization.

## CRediT authorship contribution statement

**Shuntaro Chiba:** Methodology, Software, Formal analysis, Investigation, Data Curation, Writing - Original Draft, Visualization, Funding acquisition. **Masateru Ohta:** Conceptualization, Methodology, Software, Investigation, Writing - Original Draft, Visualization, Supervision, Project administration. **Tsutomu Yamane:** Methodology, Software, Writing - Review & Editing. **Yasushi Okuno:** Writing - Review & Editing, Funding acquisition. **Mitsunori Ikeguchi:** Conceptualization, Methodology, Writing - Review & Editing, Supervision.

## Data and code availability

The descriptor calculation software intDesc-AbMut is provided at https://github.com/riken-yokohama-AI-drug/intDesc/tree/intDesc-AbMut. The dictionary-based file format converter, pdb2mol2.py, is provided at https://github.com/riken-yokohama-AI-drug/pdb2mol2. The rotamer energy calculator, rotamer_frequency.py, is provided at https://github.com/riken-yokohama-AI-drug/rotamer_frequency. In the github pages, examples for software execution are provided.

## Acknowledgements

This work was supported by JSPS KAKENHI Grant Numbers JP22K12269, the MEXT ‘Program for Promoting Research on the Supercomputer Fugaku’ (Simulation- and AI-driven next-generation medicine and drug discovery based on ‘Fugaku’, grant no. JPMXP1020230120), and the HOKUSAI BigWaterfall system and the HOKUSAI BigWaterfall2 system.

