## supplemental Information for "intDesc-AbMut: Describing and understanding how antibody mutations impact their environmental interactions^1^"

[1] **Abbreviations:** Ag/DPMAb, antigen/double-point mutant antibody; CDR, complementarity-determining region; DPM, double-point mutation; MCC, Matthews correlation coefficient; PDB, Protein Data Bank; RMSD, root mean square deviation; SPM, single-point mutation; vdW, van der Waals; WT, wild-type

**Table S1**. Definition of interactions

| Inter-action | Label | Don-or | D | A | Necessary conditions | Interaction scheme | Comment |
|---|---|---|---|---|---|---|---|
| Hydrogen bond (HB) | HB OH_O<br>HB NH_O<br>HB OH_N<br>HB NH_N | OH<br>NH<br>OH<br>NH | O<br>N<br>O<br>N | O<br>O<br>N<br>N | Dist. d(D:A) ≤ 3.2Å<br>Ang. 0°≤ ∠(H:D:A) ≤60°<br>Ang. 90°≤ ∠(H:A:An1) ≤180°<br>Ang. 60°≤ ∠(H:A:An2) ≤180° | 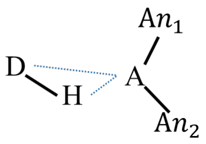 | $An_2$ is optional and can be a hydrogen. |
| Electro-static | Elec OH_O<br>Elec NH_O<br>Elec OH_N<br>Elec NH_N | OH<br>NH<br>OH<br>NH | O<br>N<br>O<br>N | O<br>O<br>N<br>N | Dist. 3.2Å ≤ d(D:A) ≤ (RvdW(D) + RvdW(A) + 1.0Å)<br>Ang. 0°≤ ∠(H:D:A) ≤60°<br>Ang. 90°≤ ∠(H:A:An1) ≤180°<br>Ang. 60°≤ ∠(H:A:An2) ≤180° | 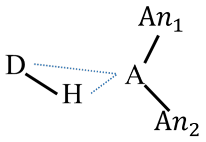 | $An_2$ is optional and can be a hydrogen. |
| CH…N | CH_N | CH | C | N | Dist. d(D:A) ≤ (RvdW(D) + RvdW(A) + 1.0Å)<br>Dist. d(D:A) ≤ d(Dn1:A)<br>Dist. d(D:A) ≤ d(D:An1)<br>Dist. d(D:A) ≤ d(D:An2)<br>Dist. d(H:A) ≤ d(D:A)<br>((Dist. d(H:A) ≤ 2.95Å) OR (Dist. (2.95Å<d(H:A) ≤ 3.10Å) AND (Ang.135° ≤ ∠(D:H:A) ≤ 180°))) | 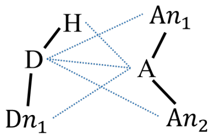 | $An_2$ is optional and can be a hydrogen. |
| CH…O | CH_O | CH | C | O | Dist. d(D:A) ≤ (RvdW(D) + RvdW(A) + 1.0Å)<br>Dist. d(D:A) ≤ d(Dn1:A)<br>Dist. d(D:A) ≤ d(D:An1)<br>Dist. d(D:A) ≤ d(D:An2)<br>Dist. d(H:A) ≤ d(D:A)<br>Dist. d(H:A) ≤ 3.22Å<br>Ang. 94.58° ≤ ∠(D:H:A) ≤ 180° | 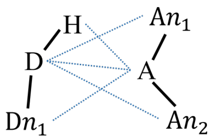 | $An_2$ is optional and can be a hydrogen. |
| CH…S | CH_S | CH | C | S | Dist. d(D:A) ≤ (RvdW(D) + RvdW(A) + 1.0Å)<br>Dist. d(D:A) ≤ d(Dn1:A)<br>Dist. d(D:A) ≤ d(D:An1)<br>Dist. d(D:A) ≤ d(D:An2)<br>Dist. d(H:A) ≤ d(D:A)<br>((Dist. d(H:A) ≤ 3.2Å) OR (Dist. (3.2Å < d(H:A) ≤ 3.3Å) AND (Ang. 135° ≤ ∠(D:H:A) ≤ 180°))) | 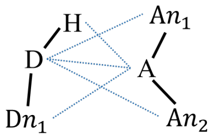 | $An_2$ is optional and can be a hydrogen. |

| | | | | | | | |
|---|---|---|---|---|---|---|---|
| OH…S | OH_S | OH | O | S | Dist. d(D:A) ≤ (RvdW(D) + RvdW(A) + 1.0Å)<br>Dist. d(D:A) ≤ d(Dn1:A)<br>Dist. d(D:A) ≤ d(D:An1)<br>Dist. d(D:A) ≤ d(D:An2)<br>Dist. d(H:A) ≤ d(D:A)<br>Ang. 90° ≤ ∠(Dn1:D:A) ≤ 180° | 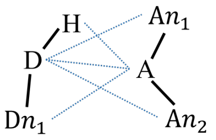 | $An_2$ is optional and can be a hydrogen. |
| NH…S | NH_S | NH | N | S | Dist. d(D:A) ≤ (RvdW(D) + RvdW(A) + 1.0Å)<br>Dist. d(D:A) ≤ d(Dn1:A)<br>Dist. d(H:A) ≤ d(D:A)<br>Dist. d(D:A) ≤ d(D:An1)<br>Dist. d(D:A) ≤ d(D:An2)<br>(Dist. (d(H:A) ≤ 3.155Å) AND (Ang. 105° ≤ ∠(D:H:A) ≤ 180°)) OR (Dist. (3.155Å< d(H:A) ≤ 3.4Å) AND (Ang. 165° ≤ ∠(D:H:A) ≤ 180°)) | 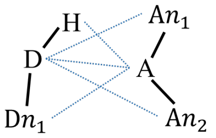 | $An_2$ is optional and can be a hydrogen. |
| SH…O | SH_O | SH | S | O | Dist. d(D:A) ≤ (RvdW(D) + RvdW(A) + 1.0Å)<br>Dist. d(D:A) ≤ d(Dn1:A)<br>Dist. d(D:A) ≤ d(D:An1)<br>Dist. d(D:A) ≤ d(D:An2)<br>Dist. d(H:A) ≤ d(D:A) | 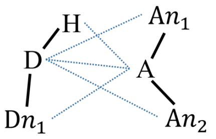 | $An_2$ is optional and can be a hydrogen. |
| SH…N | SH_N | SH | S | N | Dist. d(D:A) ≤ (RvdW(D) + RvdW(A) + 1.0Å)<br>Dist. d(D:A) ≤ d(Dn1:A)<br>Dist. d(D:A) ≤ d(D:An1)<br>Dist. d(D:A) ≤ d(D:An2)<br>Dist. d(H:A) ≤ d(D:A) | 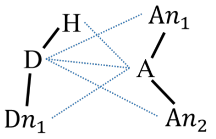 | $An_2$ is optional and can be a hydrogen. |
| SH…S | SH_S | SH | S | S | Dist. d(D:A) ≤ (RvdW(D) + RvdW(A) + 1.0Å)<br>Dist. d(D:A) ≤ d(Dn1:A)<br>Dist. d(D:A) ≤ d(D:An1)<br>Dist. d(D:A) ≤ d(D:An2)<br>Dist. d(H:A) ≤ d(D:A)<br>Ang. 0° ≤ ∠(D:H:A) ≤ 90° | 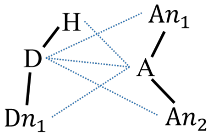 | $An_2$ is optional and can be a hydrogen. |
| $\pi-\pi$ stacking | PI_PI | Dist. d(Y:Z) ≤ (RvdW(Y) + RvdW(Z) + 1.0Å)<br>θ1 ≤ 30° if 0° ≤ θ1 ≤ 90°<br>180 - θ1 ≤ 30° if 90° < θ1 ≤ 180° | | | | 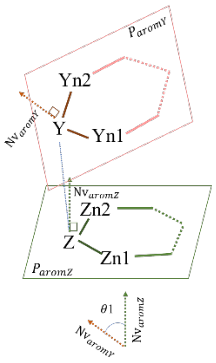 | Y, Yn1, Yn2, Z, Zn1, Zn2: aromatic. NvaromP: the normal vector of plane P. Angle θ1: the angle between NvaromY and NvaromZ |

| | | | | | | | |
|---|---|---|---|---|---|---|---|
| CH…π | CH_PI | CH | C | π | Dist. d(D:A) ≤ (RvdW(D) + RvdW(A) + 1.0Å)<br>Dist. d(D:A) ≤ d(Dn1:A)<br>Dist. d(H:A) ≤ d(D:A)<br>Dist.: d(Nrm:cn) ≤ d(cn:A)×1.4<br>(Dist. d(A:H) ≤ 3.195Å) OR ((Dist. 3.195Å<d(A:H) ≤ 3.325Å) AND (Ang. 124.455° ≤ ∠(D:H:A) ≤ 180.0°)) | 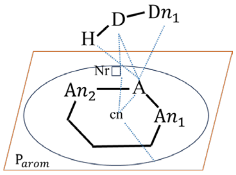 | A, $An_1$, $An_2$: Aromatic atoms |
| NH…π | NH_PI | NH | N | π | Dist. d(D:A) ≤ (RvdW(D) + RvdW(A) + 1.0Å)<br>Dist. d(D:A) ≤ d(Dn1:A)<br>Dist. d(H:A) ≤ d(D:A)<br>Dist.: d(Nrm:cn) ≤ d(cn:A) × 1.4<br>(Dist. d(A:H) ≤ 3.14Å) OR ((Dist. 3.14Å<d(A:H) ≤ 3.365Å) AND (Ang. 132.0° ≤ ∠(D:H:A) ≤ 180.0°)) | 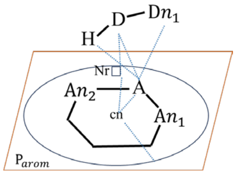 | A, $An_1$, $An_2$: Aromatic atoms |
| OH…π | OH_PI | OH | O | π | Dist. d(D:A) ≤ (RvdW(D) + RvdW(A) + 1.0Å)<br>Dist. d(D:A) ≤ d(Dn1:A)<br>Dist. d(H:A) ≤ d(D:A)<br>Dist.: d(Nr:cn) ≤ d(cn:A)×1.4<br>(Dist. d(A:H) ≤ 3.0Å) OR ((Dist. 3.0Å<d(A:H) ≤ 3.3Å) AND (Ang. 105.0° ≤ ∠(D:H:A) ≤ 180.0°)) | 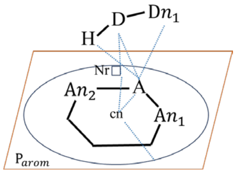 | A, $An_1$, $An_2$: Aromatic atoms |
| SH…π | SH_PI | SH | S | π | Dist. d(D:A) ≤ (RvdW(D) + RvdW(A) + 1.0Å)<br>Dist. d(D:A) ≤ d(Dn1:A)<br>Dist. d(H:A) ≤ d(D:A)<br>Dist. d(Nr:cn) ≤ d(cn:A)×1.4<br>(Dist. d(A:H) ≤ 3.2Å) OR ((Dist. 3.2Å < d(A:H) ≤ 3.33Å) AND (Ang. 120.0° ≤ ∠(D:H:A) ≤ 180.0°)) | 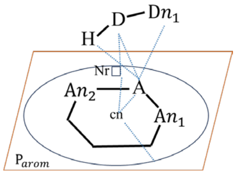 | A, $An_1$, $An_2$: Aromatic atoms |
| S…π | S_PI | S | S | π | Dist. d(D:A) ≤ (RvdW(D) + RvdW(A) + 1.0Å)<br>Dist. d(D:A) ≤ d(Dn1:A)<br>Dist. d(D:A) ≤ d(Dn2:A)<br>Dist. d(Nr:cn) ≤ d(cn:A)×1.4<br>(Ang. 30° ≤ θ1 if θ1 ≤ 90°) OR (Ang. 30° ≤ 180-θ1 if θ1>90°) | 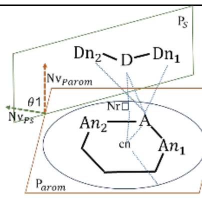 | S…π is defined if the conditions on this line or the next line are satisfied. |
| S…π | S_PI | S | S | π | Dist. d(D:A) ≤ (RvdW(D) + RvdW(A) + 1.0Å)<br>Dist. d(D:A) ≤ d(Dn1:A)<br>Dist.: d(Nr:cn) ≤ d(cn:A)×1.4<br>Dist. d(cn:D) ≤ d(cn:Dn1)<br>Ang. 120° ≤ ∠(Nr:D:Dn1) ≤ 180°<br>Dihedral angle 135° ≤ dihed(cn:Nr:D:Dn1) | 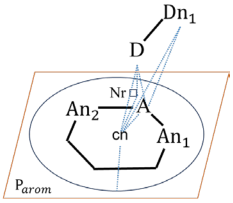 | S…π is defined if the conditions on this line or the previous line are satisfied. |

| | | | | | | | |
|---|---|---|---|---|---|---|---|
| S…O | S_O | S | S | O | Dist. d(D:A) ≤ RvdW(D) + RvdW(A) + 0.2Å<br>Dist. d(D:A) ≤ d(Dn1:A)<br>Dist. d(D:A) ≤ d(Dn2:A)<br>Dist. d(D:A) ≤ d(D:An1)<br>Dist. d(D:A) ≤ d(D:An2) | 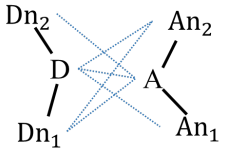  | $Dn_2$ and $An_2$ are optional and can be a hydrogen. |
| S…N | S_N | S | S | N | Dist. d(D:A)≤ RvdW(D) + RvdW(A) + 0.2Å<br>Dist. d(D:A) ≤ d(Dn1:A)<br>Dist. d(D:A) ≤ d(Dn2:A)<br>Dist. d(D:A) ≤ d(D:An1)<br>Dist. d(D:A) ≤ d(D:An2) | 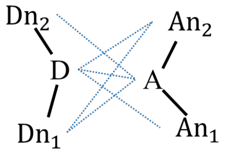  | $Dn_2$ and $An_2$ are optional and can be a hydrogen. |
| S…S | S_S | S | S | S | Dist. d(D:A)≤ RvdW(D) + RvdW(A) + 0.4Å<br>Dist. d(D:A) ≤ d(Dn1:A)<br>Dist. d(D:A) ≤ d(Dn2:A)<br>Dist. d(D:A) ≤ d(D:An1)<br>Dist. d(D:A) ≤ d(D:An2) | 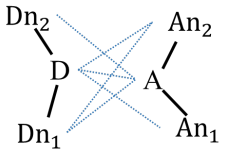  | $Dn_2$ and $An_2$ are optional and can be a hydrogen. |
| Bond dipole | DIPO | | | | $\mid \delta^+ - \delta^- \mid \geq 0.2$ in dipole 1<br>$\mid \delta^+ - \delta^- \mid \geq 0.2$ in dipole 2<br>Dist. d(Dp1+:Dp2-) ≤ (RvdW(Dp1+) + RvdW(Dp2-) + 1.0Å)<br>Ang. 0° ≤ ∠(Dp1+:Dp2-:Dp2+) ≤ 90°<br>Ang. 0° ≤ ∠(Dp2-:Dp2+:Dp1-) ≤ 90°<br>Dist. d(cn1:cn2) ≤ (RvdW(Dp1+) + RvdW(Dp2-) + 1.0Å)<br>165°≤θ1≤180° | 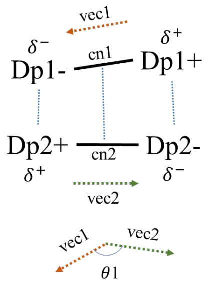  | cnX: the midpoint between DpX- and DpX+.<br>vecX: the vector from DpX+ to DpX-.<br>θ1: the angle between vec1 and vec2. |
| Orthogo-nal multi-polar interacti-on | OMulPol | | | | $\mid \delta^+ - \delta^- \mid \geq 0.2$ in dipole 1<br>$\mid \delta^+ - \delta^- \mid \geq 0.2$ in dipole 2<br>Dist. d(Dp1-:Dp2+) ≤ RvdW(Dp1-) + RvdW(Dp2+) + 0.7Å<br>Dist. d(Dp1-:Dp2+) ≤ d(Dp1+: Dp2+)<br>Ang. 75°≤ ∠(Dp2-:Dp2+:Dp1-) ≤ 105°<br>Ang. 0° ≤ ∠(Nr:Dp1-:Dp2+) ≤35°<br>Ang. 150°≤ ∠(Nr:DP1-:Dp1+) ≤180° | 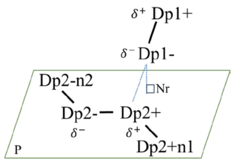  | Plane P is defined by Dp2-, Dp2+, and Dp2+n1 or Dp2-n2. |
| Metal | Fe_A(element)<br>Zn_A(element)<br>Ca_A(element)<br>Mg_A(element)<br>Ni_A(element) | | Fe<br>Zn<br>Ca<br>Mg<br>Ni | A<br>A<br>A<br>A<br>A | Dist. d(D:A) ≤ (RvdW(D) + RvdW(A) + 0.2Å)<br>Dist. d(D:A) ≤ d(D:An1)<br>Dist. d(D:A) ≤ d(D:An2) | 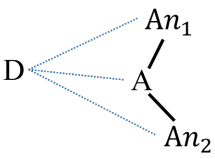  | $An_2$ is optional.<br>A(element) : the element of A. |
| Ion | Na_A(element)<br>K_A(element)<br>Cl_D(element) | | Na<br>K<br>D | A<br>A<br>Cl | Dist. d(D:A) ≤ (RvdW(D) + RvdW(A) + 1.0Å) | 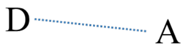  | A(element) : the element of A |
| van der Waals | vdW | | | | Dist. d(Y:Z) ≤ (RvdW(Y) + RvdW(Z) + 1.0Å) | 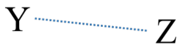  | Assigned if none of the interaction defined above. |

Label: A prefix assigned to each interaction. It is used when displaying PyMOL menus and writing descriptors. D: Donor heavy atom, A: Acceptor heavy atom, H: Hydrogen. DnN, AnN, DpN, DpNn, Y, YnN, Z, ZnN: heavy atoms. RvdW(X): VDW radius of X (Å). Black lines: Any type of bond. Blue dotted lines: Distance, Dist.: Distance, Ang.: Angle. cn: the center of the aromatic ring. P: Plane. $P_{arom}$: the plane formed by aromatic atoms. Nr: the point where the perpendicular line from the atom to the plane P intersects with P. Nv: the normal vector of the plane. $Nv_X$: the normal vector of the plane X.θ1: the angle between two normal vectors Nv.

**Table S2**. Amino acid available in pdb2mol2.py.

| Residue name | Note |
|---|---|
| GLY | |
| ALA | |
| VAL | |
| LEU | |
| ILE | |
| PRO | |
| PHE | |
| TYR | |
| TRP | |
| SER | |
| THR | |
| HID | $N_{\delta}$-protonated His |
| HIE | $N_{\varepsilon}$-protonated His |
| HIP | $N_{\delta}$- and $N_{\varepsilon}$-protonated His |
| CYS | |
| CYX | Cys with disulfide bond |
| MET | |
| LYS | |
| ARG | |
| GLN | |
| GLU | |
| ASN | |
| ASP | |

Residue names in C-terminal and N-terminal are prefixed by C and N, e.g., CGLY.

**Table S3**. PDB IDs used for training and test structure generation

| Training PDB (246 PDBs) |
|---|
| (1) 5w0d, 6wg0, 5f9o, 5dtf, 5ds8, 5dub, 2fx7, 6a77, 1ors, 1i8k, 4jfx, 3fn0, 6gku, 1yqv, 5w3p, 6p7h, 6dcw, 5ngv, 7jmp, 6b5r, 6b5s, 1ce1, 2v17, 6meh, 6pdu, 4tsb, 4tsc, 5wn9, 6aq7, 1iqd, 3mnz, 3p0y, 3sob, 6w00, 6blh, 6mtt, 1f58, 5w5z, 2ypv, 3g5y, 5vpl, 5vpg, 3rvv, 3rvw, 6mvl, 3l5x, 4g6m, 2uzi, 4hha, 6uoe, 2vxq, 6k0y, 5xku, 4dgy, 6o24, 6ulf, 6vln, 5ggv, 6d0x, 6o26, 5ob5, 2cmr, 6yio, 2adf, 7kmi, 7neh, 5vkd, 5ucb, 5uek, 6s5a, 6tgg, 6fzr, 6fzq, 6frj, 5owp, 5a2l, 5a2k, 5a2j, 1sm3, 5a2i, 6k65, 5xcq, 5tkk, 5ea0, 6z2l, 3d85, 1e4w, 6uce, 5i8c, 6ucf, 2ih3, 2hvk, 1k4c, 1r3j, 5nph, 4h88, 4ma7, 5en2, 2fd6, 4al8, 4ala, 6bfs, 5kvg, 3mxw, 6uud, 1jps, 5yy4, 5e2v, 5e2w, 1fns, 1mvu, 1h0d, 6bzy, 4r3s, 4qxt, 4qy8, 6ddr, 5lqb, 5tl5, 6j5f, 6nyq, 4m1g, 1wej, 1kir, 1kiq, 1g7m, 1g7l, 1g7j, 1g7i, 1g7h, 1a2y, 1vfb, 6b5m, 6b5n, 5hdq, 5kve, 5kvf, 6mnq, 6b0g, 3bae, 3bkj, 3eys, 5eoq, 6pxr, 1pz5, 6kx1, 6pdr, 1qkz, 3v52, 3v4u, 3uo1, 3uyr, 5myk, 6vbo, 4tuk, 4tul, 6dc8, 4h0h, 6plh, 4ojf, 4lkx, 3ley, 6x8u, 1ndg, 1nby, 1nbz, 1dqj, 2eiz, 2eks, 1ua6, 1uac, 3a6c, 3a6b, 3a67, 2dqi, 2dqe, 2dqc, 1j1x, 1j1p, 1j1o, 3d9a, 2dqd, 2dqj, 4i77, 4gag; (2) 5j56, 6zrv, 6ir1, 5vak, 4kml, 4n9o, 4nbx, 4qo1, 6yu8, 5imk, 5iml, 4dka, 6qgw; (4) 2ny4, 2ny3, 2ny2, 2nxy, 2ny1; (5) 3idg, 3drq, 2f5b, 1tjg, 1u8i; (7) 3ffd, 2qhr, 6lra; (8) 4xmp, 1op9, 3eba; (10) 3se8, 4j6r; (11) 4xvs, 4xvt; (13) 5m14, 5m15; (14) 5u3n, 5u3o; (16) 1ri8; (17) 1zv5; (19) 2xwt; (20) 2xxm; (22) 3h0t; (23) 3se9; (25) 4hpy; (26) 4nzr; (28) 4wen; (29) 5e0q; (31) 5f21; (32) 5l21; (34) 5omm; (35) 5sy8; (37) 6app; (38) 6db6; (40) 6icc; (41) 6iea; (43) 6k3m; (44) 6pec; (46) 6rtw; (47) 6u55; (49) 6vjt; (50) 6x1w |
| **Test PDB (28 PDBs)** |
| (3) 3mlr, 3uji, 3ujj, 6b0s, 4z0x, 2b1h, 4hpo; (6) 3qsk, 2p49, 2p43, 2p44; (9) 3k74, 5m2j; (12) 5fcu, 4xvj; (15) 1osp; (18) 1zvy; (21) 3go1; (24) 4gft; (27) 4orz; (30) 5e8e; (33) 5l6y; (36) 5v6m; (39) 6i2g; (42) 6ir2; (45) 6rnk; (48) 6vjn; (51) 6xzu |

In the parenthesis, CDR cluster group ID number is given. A group comprises antibodies with CDR sequence identity values > 40%.

**Table S4**. Descriptor importance

| Descriptor name | Importance (MCC decrease) | Standard deviation |
|---|---|---|
| Rot_energy | 0.2110 | 0.0114 |
| M#CH_π# | 0.1295 | 0.0106 |
| M#vdW# | 0.1294 | 0.0074 |
| M#CH_O# | 0.0726 | 0.0064 |
| M#HB_NH_O# | 0.0169 | 0.0029 |
| M#HB_OH_O# | 0.0154 | 0.0023 |
| M##S | 0.0148 | 0.0044 |
| M#Elec_OH_O# | 0.0086 | 0.0021 |
| M#π_π# | 0.0085 | 0.0025 |
| M#CH_N# | 0.0063 | 0.0019 |
| M##S## | 0.0051 | 0.0016 |
| M#NH_π# | 0.0042 | 0.0032 |
| M#S_O# | 0.0011 | 0.0006 |
| M#Dipo# | 0.0011 | 0.0011 |
| M#Elec_OH_N# | 0.0007 | 0.0006 |
| M#Elec_NH_O# | 0.0005 | 0.0019 |
| M#OMulPol# | 0.0003 | 0.0009 |
| M#HB_NH_N# | 0.0001 | 0.0004 |
| M#OH_π# | 0.0001 | 0.0003 |
| M#Elec_NH_N# | 0.0000 | 0.0000 |
| M#NH_S# | 0.0000 | 0.0000 |
| M#OH_S# | 0.0000 | 0.0000 |
| M#SH_N# | 0.0000 | 0.0000 |
| M#SH_O# | 0.0000 | 0.0000 |
| M#SH_π# | 0.0000 | 0.0000 |
| M#SH_S# | 0.0000 | 0.0000 |
| M#S_N# | 0.0000 | 0.0000 |
| M#S_π# | 0.0000 | 0.0000 |
| M#S_S# | 0.0000 | 0.0000 |
| M#HB_OH_N# | 0.0000 | 0.0001 |
| M#CH_S# | -0.0010 | 0.0012 |

**Table S5**. (a) Hyperparameters searched

| Hyperparameter | Name in XGBoost python package | Search range |
|---|---|---|
| Max depth | max_depth | 2, 3, 4, 5, 6, 8, 10 |
| Minimum sum of instance weight needed in a child | min_child_weight | 1, 3, 5, …, 31 |
| Gamma | Gamma | 0.1, 0.2, …, 0.6 |
| Subsample ratio of columns when constructing each tree | colsample_bytree | 0.3, 0.4, …, 0.9, 1 |
| Constraints of the estimation of the weights for each decision tree | max_delta_step | 0.125, 0.25, 0.5, 1, 1.5, 2, 4, 0 ("0" means no constraints.) |

(b) Hyperparameters selected

| Model | | Best hyperparameter | | | | |
|---|---|---|---|---|---|---|
| Interaction descriptor | Rotamer energy | max_depth | min_child_weight | Gamma | colsample_bytree | max_delta_step |
| + | - | 6 | 29 | 0.5 | 0.7 | 0 (No constraints) |
| + | + | 3 | 29 | 0.1 | 0.6 | 1 |

**Table S6**. (a) Predictive performance (interaction descriptors)

| Confusion matrix | | | |
|---|---|---|---|
| | | Ground truth | |
| | | Crystal-structure-like (Correct) | Non-crystal-structure-like (Incorrect) |
| Prediction | Crystal-structure-like (Correct) | 5730 | 1133 |
| | Non-crystal-structure-like (Incorrect) | 258 | 449 |
| Total | | 5988 | 1582 |
| Metric | | | |
| MCC | Accuracy | Precision | Balanced accuracy |
| 0.336 | 0.816 | 0.835 | 0.620 |

(b) Predictive performance (Rot energy + interaction descriptors)

| Confusion matrix | | | |
|---|---|---|---|
| | | Ground truth | |
| | | Crystal-structure-like (Correct) | Non-crystal-structure-like (Incorrect) |
| Prediction | Crystal-structure-like (Correct) | 5820 | 926 |
| | Non-crystal-structure-like (Incorrect) | 168 | 656 |
| Total | | 5988 | 1582 |
| Metric | | | |
| MCC | Accuracy | Precision | Balanced accuracy |
| 0.505 | 0.855 | 0.863 | 0.693 |

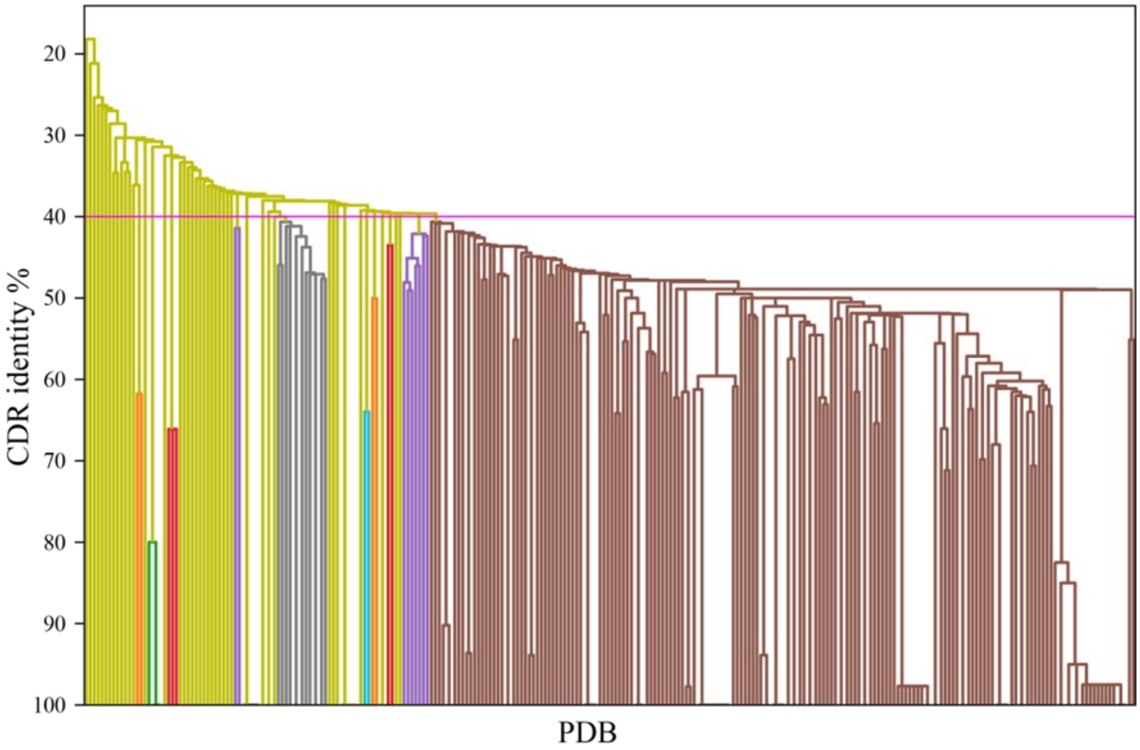


**Figure S1**. Clustering of pdb files used to derive training and test datasets. The CDR identity defined in the main text was used for the single-linkage clustering implemented in SciPy 1.5.3 [1]. The horizontal magenta line represents the 40% threshold used to define different clusters.

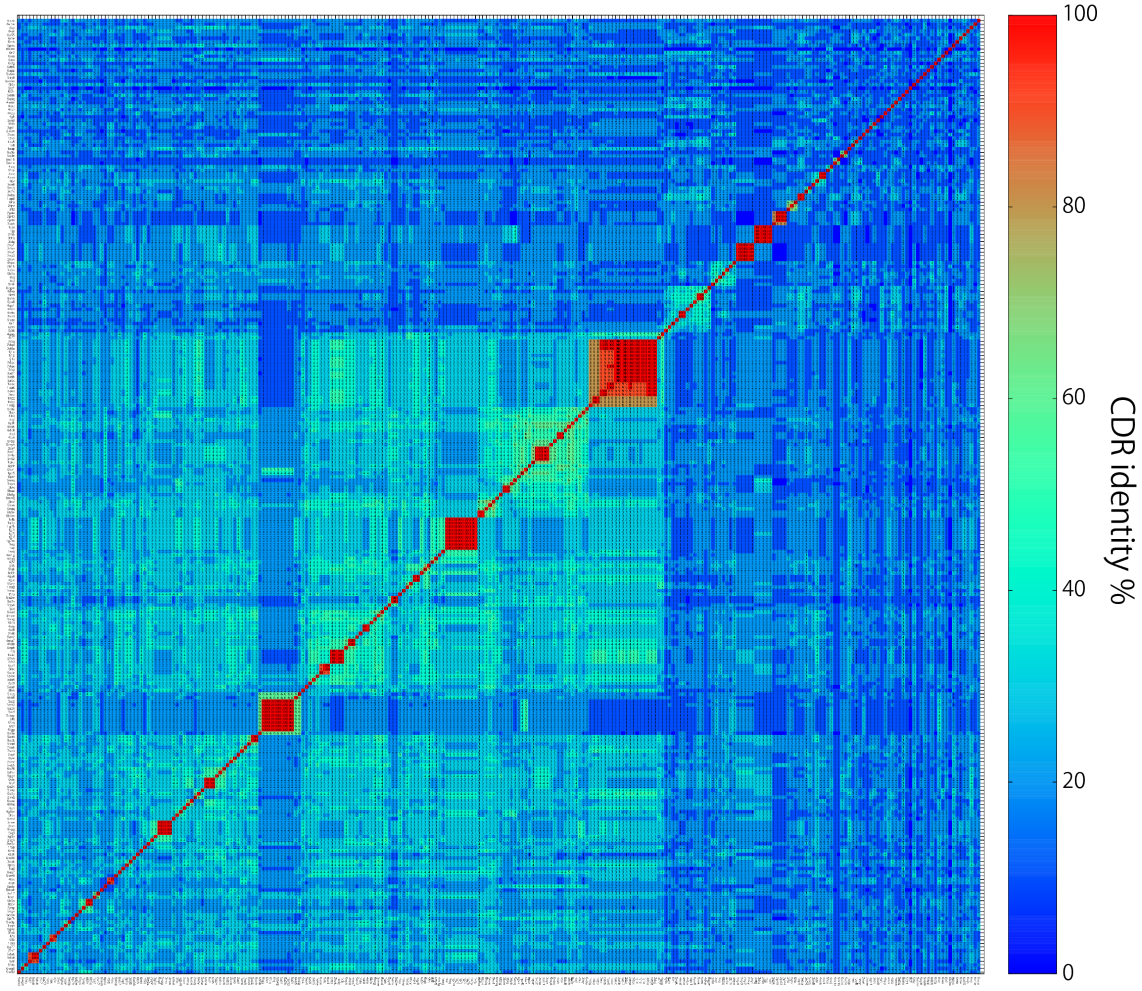


**Figure S2**. CDR identity of different PDBs sorted by cluster IDs (**Table S3**).
